# TMS-evoked EEG potentials from prefrontal and parietal cortex: reliability, site specificity, and effects of NMDA receptor blockade

**DOI:** 10.1101/480111

**Authors:** Nigel C. Rogasch, Carl Zipser, Ghazaleh Darmani, Tuomas P. Mutanen, Mana Biabani, Christoph Zrenner, Debora Desideri, Paolo Belardinelli, Florian Müller-Dahlhaus, Ulf Ziemann

## Abstract

Measuring the brain’s response to transcranial magnetic stimulation (TMS) with electroencephalography (EEG) offers a unique insight into the local cortical circuits and networks activated following stimulation, particularly in non-motor regions where less is known about TMS physiology. However, the mechanisms underlying TMS-evoked EEG potentials (TEPs) remain largely unknown. We assessed TEP reliability, site-specificity, and sensitivity to changes in excitatory neurotransmission mediated by n-methyl-d-aspartate (NMDA) receptors following stimulation of non-motor regions. In fourteen male volunteers, resting EEG and TEPs from prefrontal (PFC) and parietal (PAR) cortex were measured before and after administration of either dextromethorphan (an NMDA receptor antagonist) or placebo across two sessions separated by at least a week in a double-blinded pseudo-randomised crossover design.At baseline, TEPs showed lower within-than between-subject variability for both stimulation sites across sessions, demonstrating the reliability of non-motor TEPs within individuals. There were differences in amplitude between PFC and PAR TEPs across a wide time range (15-250 ms), however the signals were correlated after ∼80 ms, suggesting that early peaks reflect site-specific activity, whereas late peaks reflect activity patterns less dependent on the stimulated sites. TEPs were not altered following dextromethorphan compared to placebo, however low frequency resting oscillations were reduced in power. Our findings suggest that TEPs from PFC and PAR: 1) are reliable within and variable between individuals; 2) reflect stimulation site specific activity across early time points (<80 ms); and 3) are not sensitive to changes in NMDA receptor-mediated neurotransmission.

## INTRODUCTION

Transcranial magnetic stimulation (TMS) is a brain stimulation method capable of non-invasively activating cortical neurons across the scalp in humans via electromagnetic induction [Barker et al., 1985]. A single TMS pulse evokes a series of time-locked peaks and troughs in electroencephalographic (EEG) recordings of brain activity [Ilmoniemi et al., 1997], which are commonly known as TMS-evoked EEG potentials (TEPs). TEPs are reliable within and between sessions [Casarotto et al., 2010; Kerwin et al., 2018; Lioumis et al., 2009], are sensitive to changes in TMS parameters such as intensity [Casarotto et al., 2010; Rosanova et al., 2009] and pulse shape [Casula et al., 2018], and differ depending on the cortical site stimulated [Casarotto et al., 2010]. In addition, TEPs are sensitive to changes in cortical properties resulting from differing brain states, plasticity-inducing brain stimulation paradigms, and brain disorders [Rogasch and Fitzgerald, 2013]. As such, TMS-EEG is emerging as a powerful method for investigating cortical dynamics in health and disease.

Despite the recent uptake of TMS-EEG within the brain stimulation field, it remains largely unclear what physiological properties underlie the size, shape and distribution of TEPs, thereby limiting their interpretability. Current hypotheses suggest that TEPs primarily reflect fluctuations in cortical excitability resulting from excitatory and inhibitory neurotransmission at the site of stimulation, as well as the propagation of activation through cortical networks following TMS [Rogasch and Fitzgerald, 2013]. In support, pharmacological agonists of inhibitory neurotransmission mediated by fast activating γ-aminobutyric acid (GABA)-A receptors given at sub-anaesthetic doses increase the amplitude of early TEPs (e.g. N45) following motor cortex stimulation [Darmani et al., 2016; Premoli et al., 2014a], and reduce the propagation of activity following premotor and parietal cortex stimulation at anaesthetic doses [Ferrarelli et al., 2010]. Agonists of slow acting GABA-B receptors at sub-anaesthetic doses, on the other hand, increase the amplitude of latter peaks (e.g. N100) following motor cortex stimulation [Premoli et al., 2014a]. Although evidence for the sensitivity of single-pulse TEPs to inhibitory neurotransmission is growing, the effect of excitatory neurotransmission on TEPs is less clear. Several studies have linked early motor TEPs between 15-40 ms after TMS with fluctuations in cortical excitability measured via motor-evoked potentials [Bonato et al., 2006; Mäki and Ilmoniemi, 2010; Rogasch et al., 2013a]. However, TEPs following single-pulse TMS of premotor or parietal cortex are unaffected following anaesthetic doses of ketamine, an n-methyl-d-aspartate (NMDA) receptor antagonist [Sarasso et al., 2015]. To date, no studies have assessed the sensitivity of single-pulse TEPs to changes in NMDA receptor mediated neurotransmission while individuals are conscious.

The primary aim of this study was to investigate the contribution of NMDA receptor-mediated neurotransmission to the generation of TEPs following single-pulse TMS in conscious, healthy adults. We measured TEPs following prefrontal (PFC) and parietal (PAR) stimulation before and after a sub-anaesthetic dose of dextromethorphan, an NMDA receptor antagonist, or a placebo in a double-blinded pseudo-randomized crossover design. We hypothesised that early (15-40 ms) TEPs would be reduced following dextromethorphan, but not placebo. Given that TMS-EEG is a relatively new method, particularly outside of motor cortex, we also took advantage of the repeated session experimental design to: 1) characterise within- and between-subject reliability/variability in the spatiotemporal profile of TEPs; and 2) compare differences and similarities between TEPs following stimulation of different sites. As there is currently no consensus on the best way to process TMS-EEG data, we also assessed the impact of different cleaning pipelines on the study outcomes.

## MATERIALS AND METHODS

### Participants

Fifteen right-handed male participants were recruited for the study. Data from one participant was removed due to a fault in the TMS noise-masking in one condition, leaving a total of fourteen participants (mean age ± S.D. = 28.7± 5 years, range = 21-39 years). Female participants were excluded due to the possible confounding effects of the menstrual cycle on TMS-evoked cortical excitability [Smith et al., 1999]. Prior to the study, participants underwent a physical exam and were screened for contraindications to TMS [Rossi et al., 2009]. Exclusion criteria included: presence or history of neurological or psychiatric disease, current use of central nervous system active drugs, abuse of recreational drugs including nicotine or alcohol, or contraindications to dextromethorphan. The experimental procedures were approved by the local ethics committee of the Eberhard-Karls-University Medical Faculty, Tübingen (protocol 526/2014BO1), and all participants provided signed, informed consent in accordance with the latest version of the Declaration of Helsinki.

### Experimental design

Participants underwent a pseudo-randomised, placebo-controlled, double-blind cross-over experiment to assess the effects of dextromethorphan on TEPs resulting from PFC and PAR stimulation. Dextromethorphan is a non-competitive antagonist of the glutamatergic NMDA receptor, but also interacts with serotonin transporters, sigma-1 receptors, and α3β4 nicotinic acetylcholine receptors [Taylor et al., 2016]. Prior to the experimental sessions, all participants underwent a T1-weighted magnetic resonance imaging (MRI) scan of their brain for use in TMS neuronavigation and EEG electrode position digitisation (supplementary methods). Participants then attended two experimental sessions at least one week apart. During testing, participants were seated comfortably in a chair with hands resting on a pillow in their lap. Baseline measures included: systolic and diastolic blood pressure, resting motor threshold (RMT), two 4 min periods of resting EEG (eyes open and closed; measured to assess the impact of dextromethorphan on resting oscillations), and TEPs following stimulation of PFC and PAR. Following baseline measures, participants ingested either 120 mg of dextromethorphan (dosage based on previous TMS studies showing significant pharmacological effects [Wankerl et al., 2010; Ziemann et al., 1998]) or placebo (session order pseudorandomised across subjects). After 60 min, blood pressure, resting EEG, and TEP measures were repeated. 60 min was chosen based on dextromethorphan pharmacokinetics, with blood plasma levels peaking ∼60-120 min after drug ingestion [Kazis et al., 1996]. Blood pressure was measured again at the end of the experimental session.

### MRI

A T1-weighted anatomical magnetic resonance imaging (MRI) scan of the brain was acquired from each subject using a 3T MRI scanner (MAGNETOM^®^Prisma^fit^, syngo MR D13D, Siemens Healthcare GmbH. Voxel size = 1×1×1 mm^3^; FoV read = 250, FoV phase = 93.8%, TR = 2300 ms, TE = 4.18 ms, FA = 9.0°).

### EEG

EEG was recorded from 62 TMS-compatible, c-ring slit electrodes (EASYCAP, Germany) using a TMS-compatible EEG amplifier (BrainAmp DC, BrainProducts GmbH, Germany). Data from all channels were referenced to the FCz electrode online with the AFz electrode serving as the common ground. EEG signals were digitised at 5 kHz (filtering: DC-1000 Hz) and EEG electrode impedance was kept below 5 kΩ throughout the experiment. Electrode positions were digitised to each individual’s T1-weighted MR image using a frameless stereotaxic neuronavigation system (TMS Navigator, Localite GmbH, Germany). During eyes open resting EEG, participants were asked to look at a fixation cross and blink as normal. During eyes closed, participants were asked to close their eyes and avoid going to sleep.

### TMS

For TEPs, two sites were stimulated using monophasic TMS (Magstim company, UK): left superior frontal gyrus (PFC; MNI coordinates: −20, 35, 55) and left superior parietal lobule (PAR; −20, −65, 65) (see supplementary methods for TMS details). We deliberately chose sites close to the midline to minimise TMS activation of scalp/facial muscles [Mutanen et al., 2013; Rogasch et al., 2013b]. Monophasic TMS pulses (current flow = posterior-anterior in brain) were given through a figure-of-eight coil (external diameter = 90 mm) connected to a Magstim 200^2^ unit (Magstim company, UK). The TMS coil position was determined and monitored throughout the experiment using frameless stereotaxic neuronavigation co-localised to individual T1-weighted MR scans (TMS Navigator, Localite GmbH, Germany). Coil angle was positioned so that the coil handle ran perpendicular to the underlying gyrus with the handle pointing laterally. As there are currently no standardised methods for determining TMS intensity in non-motor regions, TMS intensity was set to 100% of resting motor threshold (RMT) for each site. At the beginning of each experiment, the motor hotspot for the right first dorsal interosseus (FDI) muscle was determined over left primary motor cortex as the site where slightly suprathreshold TMS pulses consistently elicited motor-evoked potentials (MEPs) in the right FDI. Electromyography was recorded using Ag-AgCl electrodes placed in a belly tendon montage over the target muscle (filter: 20-2000 Hz; sampling rate: 5 kHz). RMT (in % maximum stimulator output; MSO) was then determined as the minimum intensity to evoke at least 5 of 10 MEPs > 50 μV peak-to-peak amplitude. For the experimental conditions, 150 TMS pulses were delivered at a rate of 0.2 Hz ± 25% jitter for each site and the order of sites was randomised at each measurement point.Participants were asked to look at a fixation cross during stimulation and blink as normal.Muscle activity and excessive eye movement were monitored by an experimenter throughout the session and fed back to the participant via a tap on the shoulder if too high.

### EEG analyses

Analyses were performed in MATLAB r2017a (MathWorks Inc.) using EEGLAB (v14.1.1) [Delorme and Makeig, 2004], TESA (v0.1.0) [Rogasch et al., 2017], FieldTrip (v20170815) [Oostenveld et al., 2011], Brainstorm (v20180108) [Tadel et al., 2011], and FreeSurfer (v5.3) [Dale et al., 1999; Fischl et al., 1999] toolboxes, and custom code. All custom code is available at: (https://github.com/nigelrogasch/DXM_TMS-EEG_paper).

#### TMS-EEG

As we were interested in early TEP peaks, we developed a novel TMS-EEG cleaning pipeline including two analysis methods designed to recover early TMS-evoked activity (<45 ms) from TMS-related artifacts; the source-estimate-utilizing noise-discarding (SOUND) [Mutanen et al., 2018] algorithm and signal-space projection source-informed reconstruction (SSP-SIR) [Mutanen et al., 2016]. For each site and time point, the data were epoched around the TMS pulse (−1500 to 1500 ms), data around the TMS pulse (−2 to 6 ms) were removed and replaced with baseline data, and the average between −1000 to 1000 ms was subtracted from each epoch. Line noise was removed by fitting and subtracting a 50 Hz sine wave from the EEG time courses using linear regression, and bad channels were identified using a data-driven Wiener-estimation approach and removed [Mutanen et al., 2018]. Data were then submitted to independent component analysis (extended infomax) and components representing TMS-evoked muscle artifacts or blinks were detected using the TESA *compselect* function (default settings) and manually checked before being removed [Rogasch et al., 2017]. A high-pass filter (1 Hz, zero-phase Butterworth filter, order = 4) was applied and trials containing excessive muscle activity or movement were removed. SOUND was then applied to suppress TMS-evoked decay and other noise-related signals [Mutanen et al., 2018]. During this procedure, missing electrodes were replaced with the SOUND estimates and the data were re-referenced to average. A second round of ICA was applied, and components representing ongoing muscle activity were detected using the TESA *compselect* function (default settings) and manually checked before being removed, with special care taken not to remove components representing a mix of neural and artifactual signal. SSP-SIR was then used to suppress any remaining early TMS-related artifacts as required [Mutanen et al., 2016]. Finally, the data were downsampled (1000 Hz), low-pass filtered (100 Hz, zero-phase Butterworth filter, order = 4), re-epoched to remove possible boundary artifacts (−1000 to 1000 ms), and re-baseline corrected (−500 to −5 ms). See table S1 for number of trials, channels and components removed.

As there is currently no consensus on the best pipeline for cleaning TMS-EEG data, we re-cleaned the data using a pipeline we have used previously [Rogasch et al., 2014] (supplementary methods; table S2) and repeated the analyses to assess whether the cleaning procedure impacted the outcomes of the study.

In addition to scalp analysis, we also applied source estimation using two different methods, dipole fitting and minimum-norm estimation (MNE) [Hämäläinen and Ilmoniemi, 1994], to assess which cortical regions most likely explained the EEG scalp data. For the forward model, each individual’s T1 scan was automatically segmented using the FreeSurfer software. After visual inspections and manual corrections, the FreeSurfer output was imported to Brainstorm and the cortical surface was down sampled to 15,000 vertices. Registration between EEG and MRI was then performed by aligning the locations of EEG electrodes with the generated scalp surfaces. The head model was computed using a three-layer symmetric boundary element method via OpenMEEG [Gramfort et al., 2010], with default conductivity values (scalp = 1, skull = 0.0125 and brain= 1). For dipole fitting, each TEP topography measured at each point of time was assumed to be generated by one freely orientating current dipole located somewhere among the cortical vertices. Each of the modelled current dipoles was independently fitted to the TEP topography (least-square fit) and the location of the dipole with the best goodness-of-fit (GOF) was taken as the most likely point of TMS-evoked cortical activity [Kaukoranta et al., 1986]. For MNE, the cortical distributed sources were formed of freely orientating dipoles using the *l*^2^-MNE solution [Hämäläinen and Ilmoniemi, 1994], which was regularised with singular value decomposition using the dimensions corresponding to the 15 largest components [Mutanen et al., 2016].

#### Resting EEG

Eyes open and eyes closed resting EEG were cleaned using identical pipelines. For each condition and time point, data were downsampled (1000 Hz), bandpass (1-100 Hz) and bandstop (48-52 Hz) filtered using a zero-phase Butterworth filter (order = 4), epoched into non-overlapping 2s segments, and concatenated into a single file for each session containing eyes open and eyes closed data from pre and post drug intake measurement time points. The data were then visually inspected, and segments with excessive muscle or eye activity and noisy channels (e.g. from disconnected electrodes) were removed. Data were then submitted to the FastICA algorithm, and independent components representing blinks, eye movement, muscle activity or electrode noise were detected using the TESA *compselect* function (default settings) and manually checked before being removed. Finally, removed channels were replaced and data were re-referenced to the average of all electrodes, and separated back into individual conditions and time points. To quantify resting oscillations, data from each segment were converted into the frequency domain using a Fourier transform with a single taper Hanning window (linear trends removed; frequency resolution = 1 Hz) and then averaged across segments. See table S3 for details on number of segments, channels and components removed.

### Statistics

#### TEP variability and comparisons between sites

To assess TEP variability between sessions, absolute TEP amplitude differences averaged over electrodes and time points (15-500 ms) were calculated and compared between baseline TEPs within-subjects (dextromethorphan vs placebo session; 14 comparisons) and between-subjects (91 comparisons) using Mann-Whitney U tests. The analysis was repeated with data collapsed across time and electrodes. To assess differences in TEPs following PFC and PAR stimulation, baseline TEPs were compared between stimulation sites for each condition using cluster-based permutation statistics (cluster threshold: p<0.05 dependent t-test; cluster alpha<0.05 two-tailed; randomisation=5000; time included: 15-250 ms). To assess similarities between stimulation sites, Spearman’s correlations were performed on TEP amplitudes across electrodes (scalp) and vertices (source) for each time point, converted to z scores, and compared with baseline measures using Mann-Whitney U tests.

#### Effects of dextromethorphan on TEPs and resting oscillations

Cluster-based permutation statistics were used to compare changes in TEP amplitude and resting oscillations across time following dextromethorphan and placebo administration, and between conditions by comparing post values subtracted from pre values (cluster threshold: p<0.05 dependent t-test; cluster alpha<0.05 two-tailed; randomisation=5000). TEP analyses included a broad time range (i.e. no *a priori* assumptions about peak times; 15-250 ms), and at six peaks evident following PFC and PAR stimulation (cluster alpha<0.008; Bonferroni corrected to control the false-discovery rate testing over six peaks). For cluster-based permutation test on individual peaks, peak times were selected separately for each stimulation site by averaging baseline TEPs from each condition, and using a peak detection algorithm on the global mean field average. Data from the peaks were taken as the average of the peak ±5 ms (peaks at < 100 ms latency to TMS) or ±15 ms (peaks at > 100 ms latency to TMS). For PFC stimulation, two early peaks were not identifiable in the global mean field average, and were taken from the Fz electrode instead. Data from TEP peaks were also compared using Bayes Factor (BF) analysis to assess evidence for the null hypothesis that changes in peak amplitudes did not differ following dextromethorphan or placebo (JASP v0.8.1.2; Cauchy prior=0.07; BF^01^>3 taken as moderate evidence). For Bayes Factor analysis, data from the six highest amplitude electrodes were averaged for each peak and post values subtracted from the pre values to create a single change score for each condition. For resting oscillations, data were averaged into five canonical oscillation bands prior to cluster-based analysis: delta (1-3 Hz); theta (4-7 Hz); alpha (8-12 Hz); beta (13-29 Hz); and gamma (30-45 Hz) (cluster alpha<0.01; Bonferroni corrected to control the false-discovery rate testing over five bands).

## RESULTS

All experimental procedures were generally well tolerated, with several individuals reporting mild dizziness and one individual nausea following dextromethorphan. These side effects did not affect the subjects’ capacity to fully comply with study requirements. There was no difference in RMT at baseline between drug conditions (dextromethorphan=48.4 ± 8% maximum stimulator output, MSO; placebo=47.8 ± 8% MSO; p=0.09). Changes in blood pressure did not differ between conditions (supplementary materials).

### Within- and between-subject TEP variability

We first compared within- and between-subject differences in baseline TEPs across conditions to assess TEP reliability/variability. Differences in TEPs across sessions within individuals were lower than differences between individuals for both stimulation sites (PFC, p=4.7×10^−6^; PAR, p=1.1×10^−4^; fig. 1A,B). Across space, lower within-subject than between-subject TEP differences were observed across the majority of electrodes for both stimulation sites (fig. 1C,D), whereas across time, lower within-subject TEP differences were stronger between ∼30-513 ms following PFC stimulation and ∼25-323 ms following PAR stimulation (fig. 1E,F). These findings suggest that the spatiotemporal profile of TEPs are reliable across sessions within individuals, but show variability between individuals (see fig. 2 for an example in 5 subjects).

**Figure 1:**
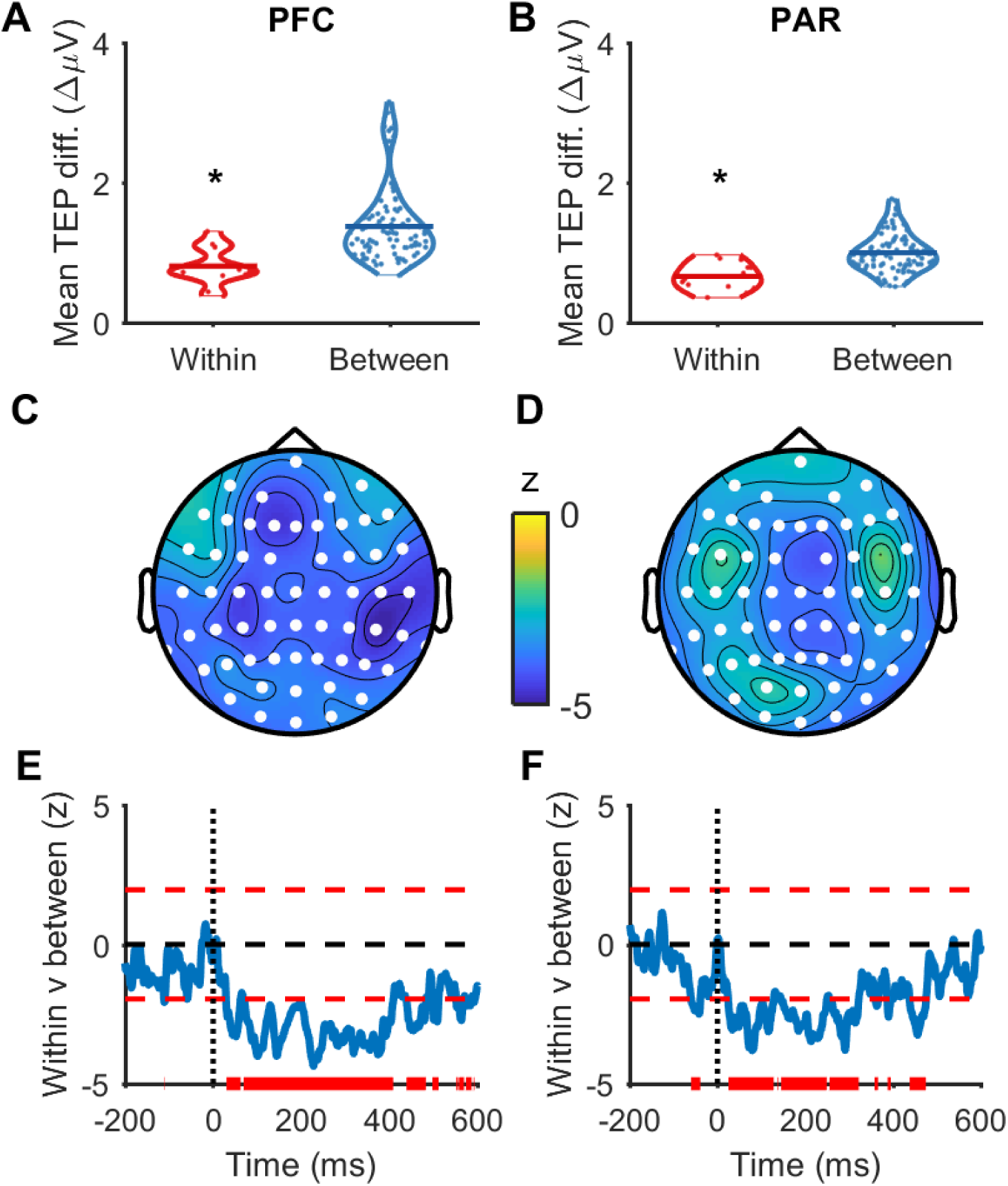
Within- and between-subject variability in baseline TEPs. A-B) Mean absolute differences in baseline TEPs (15-500 ms, all electrodes) between the dextromethorphan and placebo condition within- and between-subjects following prefrontal cortex (PFC; A) and parietal cortex (PAR; B) stimulation. * indicates p<0.05 (Mann-Whitney U test). C-D) Topoplots displaying z-scores (Mann-Whitney U tests) comparing within- and between-subject baseline TEP differences at individual electrodes (averaged across time between 15-500 ms) following PFC (C) and PAR (D) stimulation. Negative z-scores indicate within-subject TEP differences are less than between-subject TEP differences. White dots indicate p<0.05. E-F) Z-scores (Mann-Whitney U tests) comparing within- and between-subject TEP differences at individual time points (averaged across all electrodes) following PFC (E) and PAR (F) stimulation. Dotted black lines indicate the time of the TMS pulse. Dashed red lines indicate z = ±1.96. Solid red lines indicate p<0.05.

**Figure 2:**
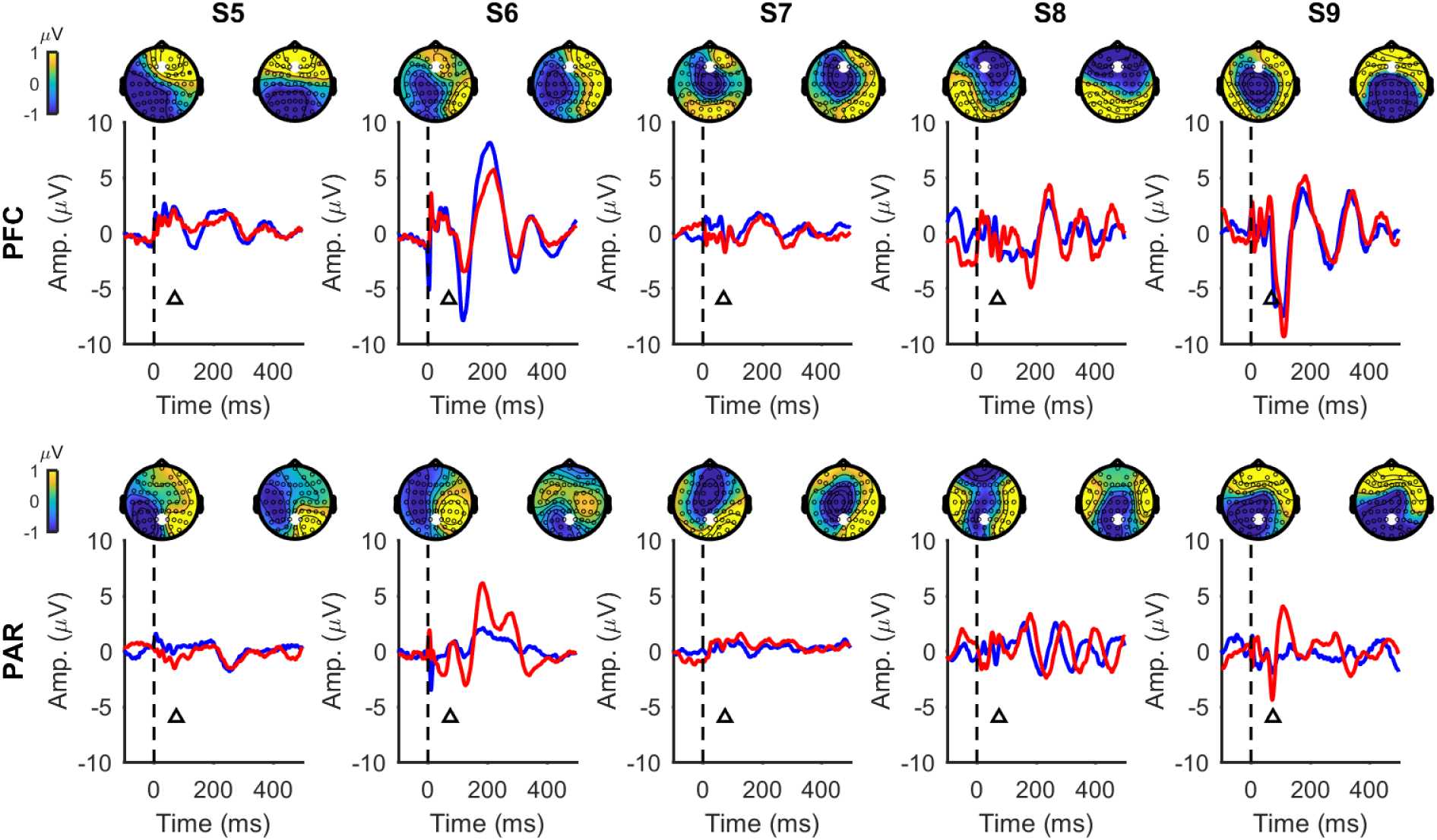
Example of within- and between-subject variability in individual participants. The top row shows baseline TEPs from the dextromethorphan (blue lines, left topoplots) and placebo (red lines, right topoplots) sessions following PFC stimulation, and the bottom row following PAR stimulation, in five example participants (S5-S9). TEP line plots are taken from an electrode near the site of stimulation (indicated with white dot on topoplots; Fz for PFC stimulation; Pz for PAR stimulation), and TEP topoplots for a representative point in time (indicated with triangles in line plots; 70 ms for PFC; 75 ms for PAR). Both the shape and spatial distribution of the baseline TEPs are more similar within-subjects than they are between-subjects.

### Baseline TEPs following PFC and PAR stimulation

Next we assessed the differences and similarities between TEPs following stimulation of different sites. As TEPs were reliable within individuals between sessions, we averaged across baseline conditions to maximise TEP signal strength. When comparing across a broad time window (15-250 ms), TEPs following PFC stimulation differed in amplitude compared with PAR stimulation across all time points (fig. 3). Despite the amplitude differences, the spatial distribution of TEPs were highly correlated between stimulation sites after ∼83 ms (fig. 4A), suggesting that later peaks may represent similar underlying cortical sources regardless of the stimulated sites.

**Figure 3:**
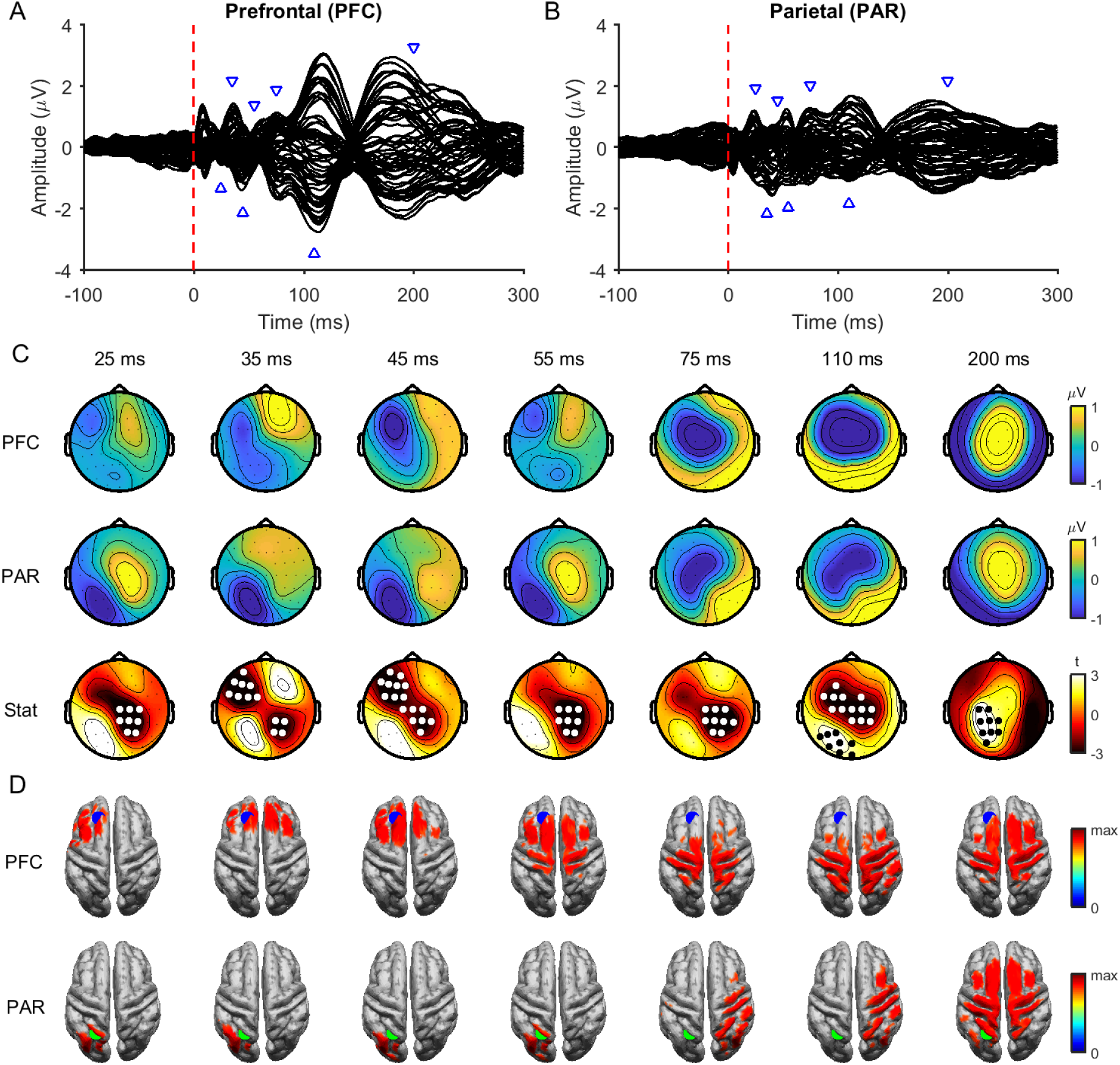
Comparison of baseline TEPs between stimulation sites. Butterfly plots of grand average TEPs across all individuals following prefrontal (PFC; A) and parietal cortex (PAR; B) stimulation at baseline (averaged across conditions). The red dashed line represents the timing of the TMS pulse and the blue triangles the latencies plotted in C and D. C) Topoplots showing the grand average amplitude of TEPs at different time points following PFC (top row), and PAR stimulation (middle row). The bottom row shows t-statistics comparing the amplitude of PFC and PAR stimulation. White and black dots indicate significant negative and positive clusters (cluster-based permutation tests on 15-250 ms; 2 positive clusters [p=0.040, 81-142 ms; p=0.006, 148-250 ms]; 1 negative cluster [p=0.002, 15-192 ms]). D) Minimum-norm estimate source maps averaged across participants showing peak activity at each time point in C following PFC (top row) and PAR (bottom row) stimulation. Activity has been thresholded to 85% of maximum activity at each time point. The blue dot represents the target for PFC stimulation and the green dot the target for PAR stimulation.

**Figure 4:**
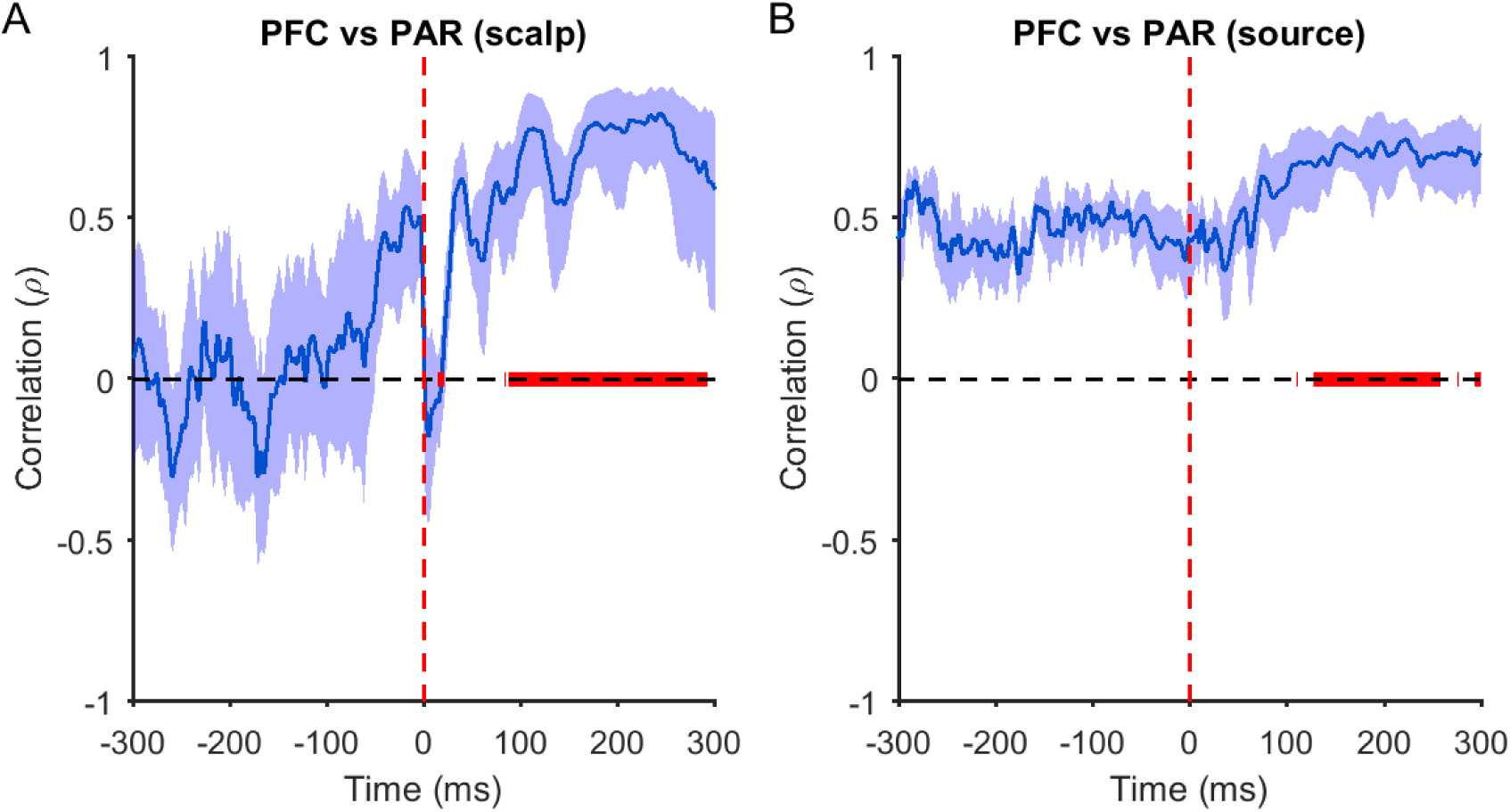
Spatial correlations between prefrontal (PFC) and parietal (PAR) TEPs. Spearman correlations comparing the relationship between PFC and PAR TEPs at the scalp (A) and source (B) level for each time point. The thick blue line represents the mean rho values across individuals, and the shaded bars the 95% confidence intervals. The thick red line indicates post stimulation time points where correlations are greater than at equivalent pre stimulation time points (p<0.05; Mann-Whitney U test). Note that rho values were converted to z for statistics, then back to rho for plotting.

To further explore the origin of early and late TEPs, we applied two different source estimation methods: dipole fitting and MNE. For early peaks, the location of the best fitting dipole tended to be closer to the site of stimulation compared to the non-target site (e.g. the PAR when the PFC was stimulated and *vice versa*; table 1). In contrast, the dipole locations corresponding to late peaks were closer to the PAR target regardless of stimulation site. For MNE, estimated source distributions were located close to the site of stimulation for early peaks (25-55 ms; fig. 3D), showed some overlap between stimulation sites for middle peaks (75,110 ms), and were similar for late peaks (200 ms). Similar to the scalp data, MNE spatial distributions were highly correlated between PFC and PAR TEPs from ∼129 ms to ∼259 ms (fig. 4B). Taken together, these findings suggest that early TEP peaks reflect neural activation specific to the site of stimulation, whereas late peaks reflect common activation patterns, which differ in amplitude between stimulation sites.

**Table 1:** Distance from TMS target sites to best-fitting dipoles at baseline.

|  | Distance<br>from target<br>(mm) | Distance from<br>non-target<br>(mm) | Goodness of<br>fit (GoF) | p-value |
| --- | --- | --- | --- | --- |
| PFC (15-45 ms) | 60<br>[18-133] | 73<br>[40-103] | 0.93<br>[0.81-0.99] | 0.135 |
| PFC (95-125 ms) | 80<br>[24-129] | 59<br>[20-110] | 0.88<br>[0.68-0.99] | 0.077 |
| PFC (175-205 ms) | 80<br>[37-123] | **51**<br>**[21-122]** | 0.84<br>[0.69-0.99] | 0.003 |
| PAR (15-45 ms) | **49**<br>**[23-91]** | 97<br>[68-130] | 0.93<br>[0.69-0.99] | 4.7x10<sup>-5</sup> |
| PAR (95-125 ms) | **52**<br>**[22-94]** | 87<br>[51-110] | 0.90<br>[0.79-0.97] | 1.5x10<sup>-4</sup> |
| PAR (175-205 ms) | **44**<br>**[19-85]** | 82<br>[42-130] | 0.87<br>[0.72-0.99] | 0.001 |

**NB:** Values in column 1-3 represent the mean [range]. Bold numbers indicate which site was closest to the best fitting dipole (target vs. non-target; $p < 0.05$ , Mann-Whitney U test). PFC, prefrontal cortex; PAR, parietal cortex.

### Effect of dextromethorphan on TEPs

We next assessed whether dextromethorphan altered TEP amplitude. We could not find any differences in TEP amplitudes across time following either dextromethorphan or placebo for PFC stimulation (all p>0.05), whereas there was a change in PAR TEP amplitude following dextromethorphan (positive cluster, p=0.006, 126-207 ms; negative cluster, p=0.0132, 125-201 ms), but not following placebo (p>0.05). However, these changes were not aligned to TEP peaks (fig. S1) and we could not find any difference between conditions when directly comparing the change in TEP amplitudes following dextromethorphan and placebo for either stimulation site (all p>0.05; 15-250 ms; fig. 5), suggesting the changes observed following dextromethorphan with PAR stimulation were not robust. To ensure that the size of latter clusters was not biasing the analysis against smaller earlier clusters, we reran the analysis averaging across shorter time windows capturing the main TEP peaks, but could not detect any differences across time or between conditions (all p>0.05; Bonferroni corrected; fig. 6). We then ran Bayesian t-tests over ROIs for each peak (determined from baseline data) to assess evidence for the null hypothesis that changes in TEP amplitudes did not differ between conditions. For all comparisons, the BF_01_ was between 1-4, providing weak/moderate evidence that changes in TEP peak amplitude did not differ between dextromethorphan and placebo (table 2).

**Table 2:** Bayes factors comparing the change in TEP peak amplitude following dextromethorphan (DXM) vs. placebo (PBO).

| TEP peaks | DXM vs PBO |  |
| --- | --- | --- |
|  | PFC (BF<sub>01</sub>) | PAR (BF<sub>01</sub>) |
| 33, 25 | **3.0** | 1.3 |
| 43, 41 | 2.5 | **3.7** |
| 60, 54 | **3.6** | 2.6 |
| 77, 73 | **3.7** | **3.6** |
| 115, 112 | **3.6** | 1.9 |
| 184, 194 | **3.4** | **3.0** |

**NB:** Values in column one represent the mean TEP peak latency for prefrontal cortex (PFC) and parietal cortex (PAR) stimulation respectively. Bold numbers indicate moderate evidence for no difference between conditions.

**Figure 5:**
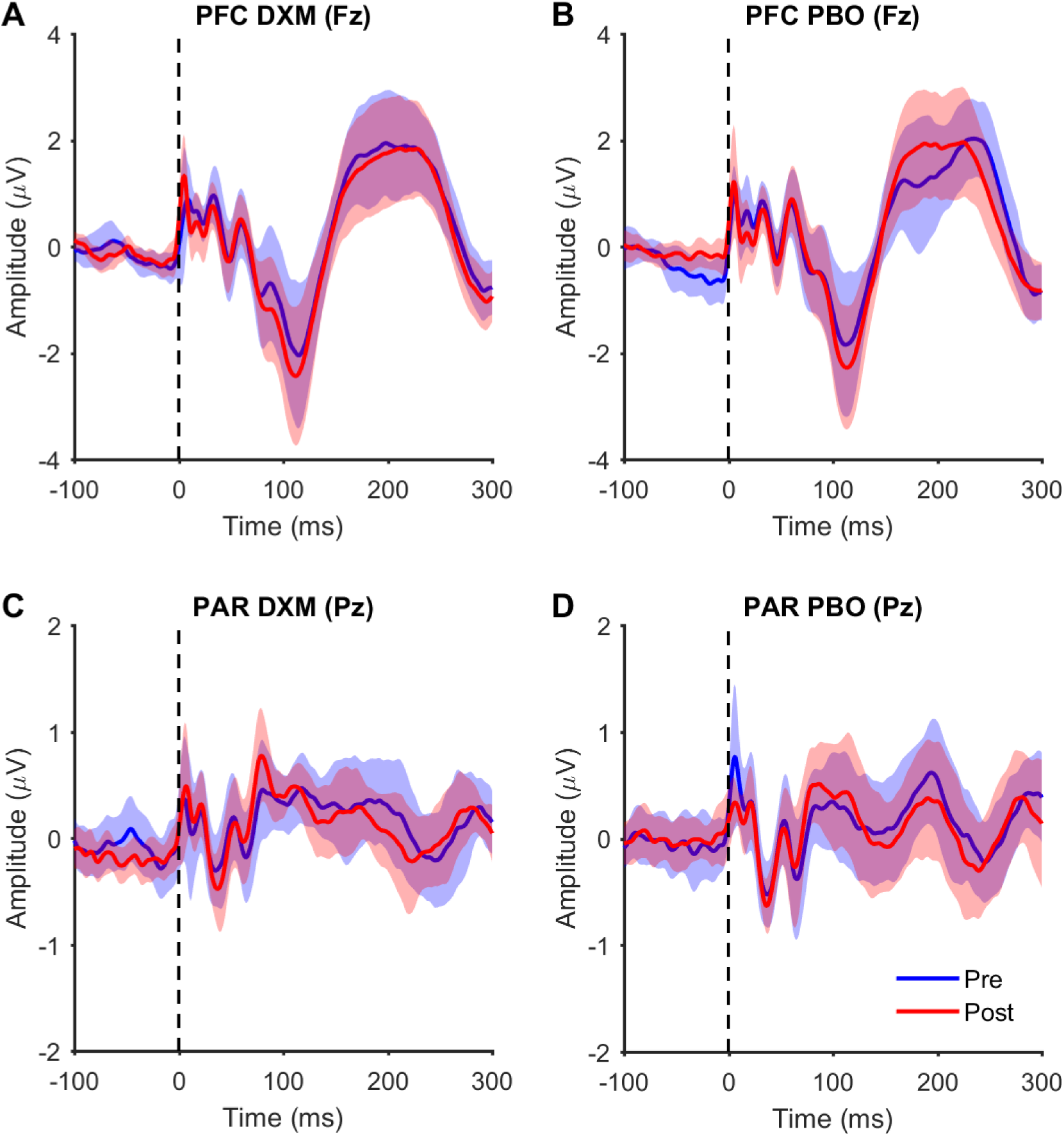
TEPs from single electrodes following dextromethorphan (DXM) and placebo (PBO). A-B) TEPs measured from the Fz electrode following prefrontal cortex (PFC) stimulation pre and post dextromethorphan and placebo administration. C-D) TEPs measured from the Pz electrode pre and post dextromethorphan and placebo administration. Thick coloured lines represent the group mean and shaded colour lines represent 95% confidence intervals.

**Figure 6:**
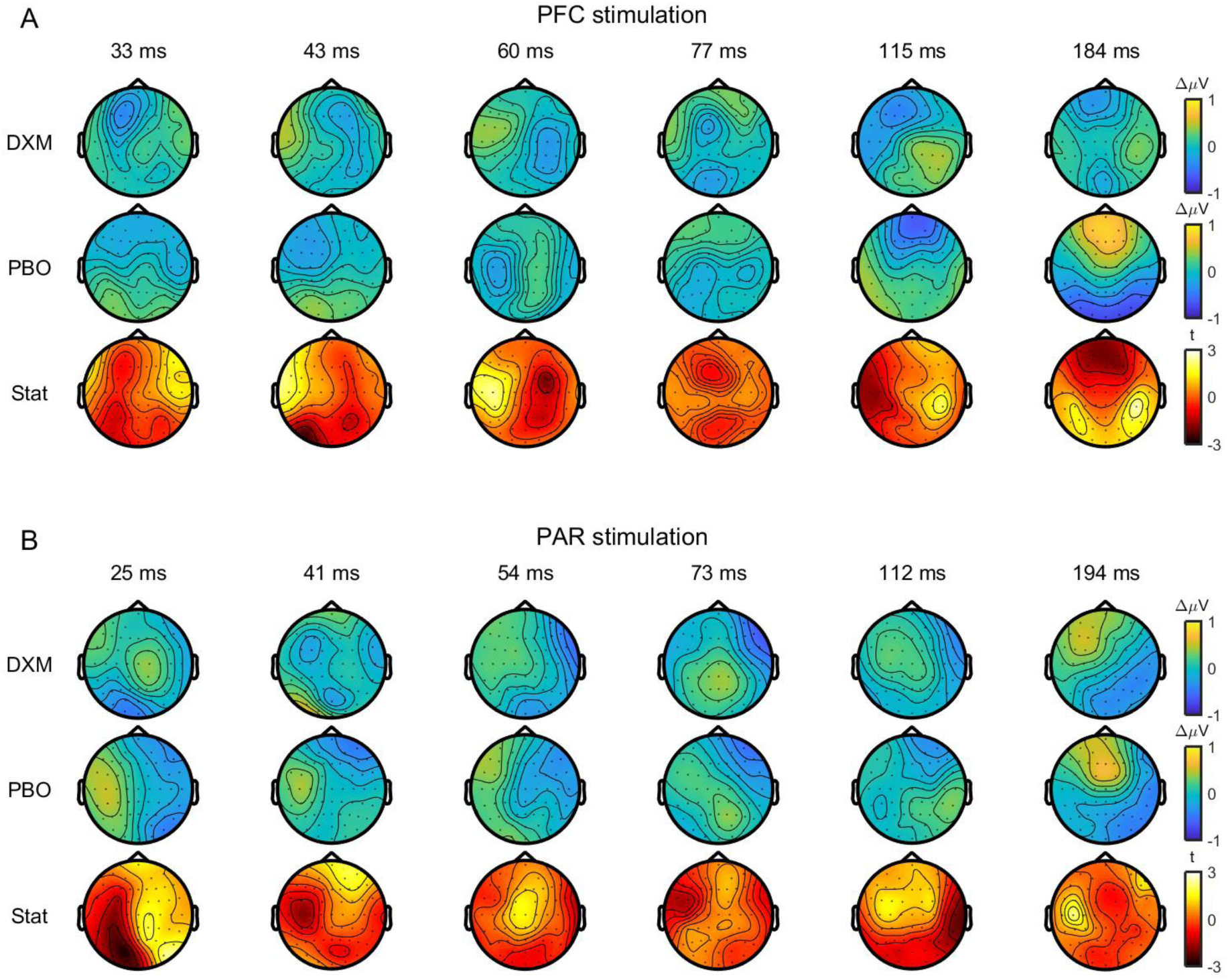
Comparison of changes in TEPs following dextromethorphan (DXM) and placebo (PBO). Topoplots showing changes in TEP amplitude at peak latencies following prefrontal (PFC; A) and parietal cortex (PAR; B) stimulation after dextromethorphan (top row) and placebo (middle row). Topoplots showing t-statistics (within-subject t-tests) comparing TEP changes between dextromethorphan and placebo are shown on the bottom row. No significant differences were observed between conditions (cluster-based permutation tests).

### Effect of processing pipeline on TEP results

As we used a novel TEP cleaning pipeline, we reran all of the analyses using a more conventional pipeline with two rounds of ICA [Rogasch et al., 2014; Rogasch et al., 2017]. As with pipeline one, we found low within-subject TEP differences between sessions, differences and similarities in amplitude between stimulation sites, and non-significant effects of dextromethorphan on TEPs using pipeline two (figs. S2-S7; tables S4-S5), indicating that the choice of cleaning pipeline does not impact the main conclusions of the study.

### Effect of dextromethorphan on resting oscillations

In addition to TEPs, we also assessed whether dextromethorphan altered resting oscillations. We could not detect any differences in resting oscillations at baseline between sessions (all p>0.05), suggesting that the spatio-spectral profile of oscillations was stable across sessions within individuals. Delta oscillatory power was reduced following dextromethorphan in the eyes open (p=0.002) and eyes closed (p=0.009) conditions, whereas beta oscillatory power was reduced following placebo in the eyes closed condition only (p=2.0×10^−4^). When comparing conditions, reductions in delta power tended to be larger following dextromethorphan than placebo for eyes open (p=0.013; fig. 7A), although this did not survive correction for multiple comparisons, whereas a reduction in theta power was larger following dextromethorphan than placebo for the eyes closed condition (p=0.009; Bonferroni-corrected; fig. 7B). We could not detect differences in oscillatory power changes between dextromethorphan and placebo for any other frequency band (all p>0.05). Taken together, these findings suggest that dextromethorphan reduces power in low frequency oscillations (delta and theta) during resting states.

**Figure 7:**
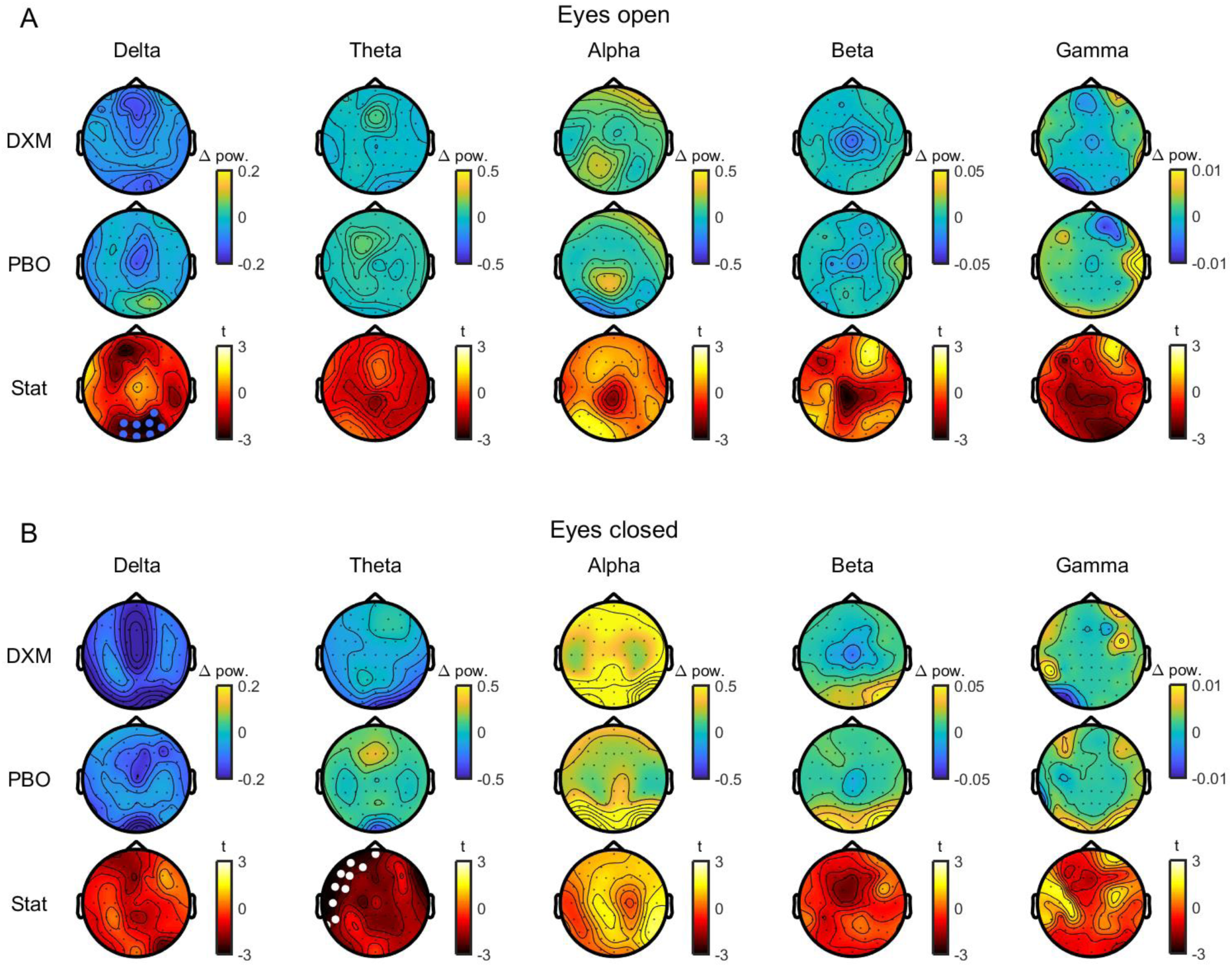
Comparison of changes in resting oscillations following dextromethorphan (DXM) and placebo (PBO). Topoplots showing changes in oscillatory power in different frequency bands during eyes open (A) and eyes closed (B) resting conditions following dextromethorphan (top row) and placebo (middle row). Topoplots showing t-statistics (within-subject t-tests) comparing power changes between dextromethorphan and placebo are shown on the bottom row. White dots indicate significant clusters with Bonferroni correction and blue dots uncorrected clusters.

## DISCUSSION

TEPs offer unique insight into the effects of TMS on local cortical circuits and networks, however the precise mechanisms reflected by TEPs remain largely unclear. In the current study, we have shown that TEPs from PFC and PAR stimulation are reliable within-individuals, but are variable between-individuals. When comparing TEPs from different stimulation sites, early TEPs (<50 ms) are localised to regions close to the site of stimulation, whereas late peaks (>80 ms) showed common activation patterns, independent of the stimulated sites. We also provide weak/moderate evidence that TEPs are not altered by dextromethorphan, suggesting that TMS-evoked EEG responses following single-pulses applied to PFC and PAR are insensitive to changes in NMDA receptor mediated neurotransmission. Our findings confirm the reliability of TEPs for assessing the cortical response to TMS in regions outside the motor cortex, and provide a deeper understanding of the physiological mechanisms reflected by TEPs elicited by prefrontal and parietal cortex stimulation.

### Reliability and variability of TEPs

TEPs are generally considered highly reliable within-individuals over short (e.g. hours) to moderate (e.g. weeks) time periods. Indeed, high test-retest reliability has been demonstrated for single-pulse TEPs following stimulation of motor [Lioumis et al., 2009; Premoli et al., 2014b], premotor [Casarotto et al., 2010], dorsolateral prefrontal [Kerwin et al., 2018; Lioumis et al., 2009], superior parietal [Casarotto et al., 2010], and visual cortices [Casarotto et al., 2010] using a variety of different metrics. We further confirm the reliability of TEPs following stimulation of superior frontal and superior parietal cortex using a simple metric which takes into account the entire spatio-temporal TEP profile by comparing the mean absolute difference in TEP amplitude within- and between-individuals across sessions (i.e. weeks). In addition to demonstrating within-individual reliability, this finding also suggests that TEPs are variable between individuals (e.g., fig. 2). The sources of this variability are likely multi-faceted, and could reflect individual differences in cortical physiology and anatomy (e.g. balance of excitation/inhibition, shape of gyri/sulci) and/or experimental factors (e.g., success of sensory masking, relative stimulation intensity). Understanding what contributes to this between-individual variability in TEP profile will likely provide important insights into how individual physiology shapes the local and global cortical response to TMS from regions outside of the motor cortex.

### Dependence of TEPs on stimulation site

Studies directly comparing TEPs following stimulation of different cortical sites have shown both differences and similarities in the local response profile and the cortical networks activated by TMS. For instance, the local oscillatory profile following TMS appears to differ along an anterior-posterior gradient, with frontal sites oscillating at higher frequencies than parietal and occipital sites [Rosanova et al., 2009]. Furthermore, stimulation of motor cortex results in larger TEPs than non-motor regions [Kähkönen et al., 2004], with a unique oscillatory profile [Fecchio et al., 2017]. The broader cortical networks activated following TMS also differ depending on the stimulation site, even within stimulation of functionally-related regions [Garcia et al., 2011].

Despite the differences in TEPs following stimulation of different cortical sites, several studies have reported similarities in TEPs regardless of the target site, especially at periods ∼100 ms, and ∼200 ms following stimulation [Du et al., 2017]. These periods coincide with auditory-evoked potentials resulting from the TMS clicking noise, and bone-conducted sensory responses from coil vibration [Nikouline et al., 1999]. To minimise sensory contamination, noise-masking is typically provided during stimulation (e.g. white noise played through headphones) and/or foam is placed under the coil to minimise vibration [Massimini et al., 2005]. Even with such measures, several recent studies have reported that TEPs are highly correlated with control conditions (e.g. TMS of the shoulder or electrical stimulation of the scalp) [Conde et al., 2018; Gordon et al., 2018; Herring et al., 2015].

In the current study, we applied auditory masking, and stimulated sites close to the midline to minimise sensation resulting from the activation of scalp muscles with TMS. We found differences in TEP amplitude following stimulation of PFC and PAR at the scalp level across a broad time range (15-250 ms). However the spatial distribution of the TEPs were highly correlated between sites from ∼80 ms onwards. Source estimation using two different methods (dipole fitting and MNE) suggested that the early TEP response (15-55 ms) reflected activity from regions close to the site of stimulation, whereas late TEP responses reflected activity from partially or fully overlapping central regions regardless of stimulation site. One possible explanation for this finding is that part of the late TEP response reflects indirect activation of the cortex from sensory input, regardless of the efforts to minimise TMS-evoked sensation and audition. Another possibility for explaining similarities in spatial distribution of late TEPs in the present study is that areas of the fronto-parietal network were stimulated potentially leading to common network activation at late time points.

### Effects of dextromethorphan on TEPs

Pharmacological studies targeting inhibitory receptors have provided evidence that certain TEP peaks around 45 and 100 ms are sensitive to changes in GABAergic neurotransmission [Darmani et al., 2016; Premoli et al., 2014a], whereas peaks at 30 ms, 45 ms and 180 ms are sensitive to anti-epileptic drugs targeting voltage-gated sodium channels [Darmani et al., 2018; Premoli et al., 2017a]. However, the sensitivity of TEPs to changes in excitatory neurotransmission is less clear. Several lines of indirect evidence suggest that early TEP peaks between 15 to 40 ms may reflect excitatory neurotransmission. First, excitatory postsynaptic potentials generated by NMDA receptor activation peak at ∼15-40 ms in rodents following electrical stimulation of the neocortex [Sutor and Hablitz, 1989], latencies which are similar to early TEP peaks. Second, the amplitude of early TEP peaks in motor cortex (N15, P30) correlate with fluctuations in MEP amplitude (which reflect activation of the corticomotoneuronal system) [Mäki and Ilmoniemi, 2010], and show similar changes with TMS intensity [Komssi et al., 2004], coil angle [Bonato et al., 2006; Komssi et al., 2004], and paired pulse paradigms [Rogasch et al., 2013a] to MEPs. Collectively, this body of evidence has led to the hypothesis that early TEP peaks may reflect excitatory postsynaptic potentials following TMS, possibly mediated by NMDA receptors [Rogasch and Fitzgerald, 2013].

We could not find any evidence that changes in TEPs differed following administration of the NMDA receptor antagonist dextromethorphan compared to placebo at any time point following stimulation of either site. The lack of change in TEPs following dextromethorphan was not impacted by our choice of statistical approach or the TEP processing pipeline. Although our sample was relatively small (n=14), Bayes factor analysis provided moderate evidence for the null hypothesis in 8 of the 12 TEP peaks tested across sites, and weak evidence in the other peaks, suggesting that we were adequately powered to test our hypothesis. In line with our findings, TEPs following single-pulse TMS to premotor and parietal cortex are largely unaffected by anaesthetic doses of ketamine [Sarasso et al., 2015], another NMDA receptor antagonist, suggesting that single-pulse TEPs are insensitive to changes in NMDA receptor mediated neurotransmission. As NMDA receptors are dependent both on glutamatergic binding and depolarisation of the postsynaptic neuron, it is possible that a single TMS pulse is not sufficient to open NMDA receptors. Instead, paired-pulse TMS-EEG paradigms at intervals between 10-40 ms may be required to observe NMDA receptor mediated neurotransmission [Cash et al., 2017], similar to intracortical facilitation paradigms measured with MEPs in motor cortex [Ziemann et al., 1998]. Alternatively, early TEPs may reflect neurotransmission mediated by other ionotropic glutamate receptors, such as AMPA receptors, which requires further investigation.

### Effects of dextromethorphan on resting oscillations

Sub-anaesthetic doses of NMDA receptor antagonists, such as ketamine, have been reported to reduce power in delta, posterior theta and alpha oscillations, and increase frontal theta and gamma oscillations in human resting EEG [de la Salle et al., 2016] and magnetoencephalographic [Muthukumaraswamy et al., 2015] recordings. We partially replicate these findings with dextromethorphan, showing reduced delta and theta oscillation power compared to placebo, however no changes in alpha or gamma oscillations. The reasons why dextromethorphan did not increase gamma oscillation power is unclear, although similar findings have been reported in animal models [Sagratella et al., 1992]. Our findings add to the growing body of evidence demonstrating an important role for NMDA receptors in low frequency oscillations.

### Limitations of the study

A potential limitation of the current study is the dose of dextromethorphan provided (120 mg) is lower than that required to produce hallucinations and cognitive impairment [Carter et al., 2013], which are hallmarks of the effects of ketamine., However, we did observe modulation of low frequency resting oscillations similar to those observed with ketamine, and dextromethorphan at similar doses blocks paired pulse and plasticity effects mediated by NMDA receptors in other TMS paradigms [Wankerl et al., 2010; Ziemann et al., 1998], suggesting the dose here was adequate. Another potential limitation is that we only tested TEPs at one intensity. The effect of certain drugs can impact TEPs in a way which is dependent on stimulation intensity [Premoli et al., 2017b]. As such, future studies assessing drug effects on TEPs should take into account a range of stimulation intensities.

## CONCLUSIONS

Our findings provide evidence that single-pulse TEPs following stimulation of prefrontal and parietal cortex in conscious humans are not sensitive to changes in excitatory neurotransmission following NMDA receptor antagonism with dextromethorphan, at least at the dose tested. However, TEPs from these cortical regions are reliable within-individuals, and the early TEP peaks provide information specific to the site of stimulation, whereas late TEPs reflect activity less dependent on the stimulated sites. Future work using pharmacological agents targeting different excitatory and inhibitory receptor types is required to disentangle the physiological mechanisms contributing to early TEPs following TMS, and to test if these pharmacological effects are different when stimulating different cortical sites.

## ACKNOWLEDGMENTS

The authors wish to thank Dr. Ben D. Fulcher for advice on analysis. This work was supported by the National Health and Medical Research Council of Australia [grant number: GNT1072057], and the Deutscher Akademischer Austauschdienst (German Academic Exchange Service).

