## Supplementary material for "TMS-evoked EEG potentials from prefrontal and parietal cortex: reliability, site specificity, and effects of NMDA receptor blockade"

**SUPPLEMENTARY METHODS**

*TMS-EEG cleaning pipeline two:* For each site, data were epoched around the TMS pulse (-1500 to 1500 ms) and baseline corrected (-1000 to -10 ms). Data around the TMS pulse were removed (-2 to 10 ms) and replaced by cubic interpolation prior to downsampling (1000 Hz), and then pre and post drug intake measurement time points concatenated so that independent component analysis (ICA) was applied equally across time. The data were then visually inspected, and epochs with excessive muscle or eye activity were removed, as were electrodes with excessive noise (e.g. from contact with the TMS coil). Data were then submitted to two rounds of FastICA [Hyvärinen and Oja, 2000]. In the first round, independent components representing TMS-evoked muscle or large decay artifacts were detected using the TESA compselect function (default settings) and manually checked before being removed [Rogasch et al., 2014; Rogasch et al., 2017]. Data were then bandpass (1-100 Hz) and bandstop (48-52 Hz) filtered using a zero-phase butterworth filter (order = 4) prior to the second round of FastICA, in which components representing blinks, eye movement, muscle activity or electrode noise were detected with TESA and manually removed. Finally, missing electrodes were replaced using spherical interpolation before data were re-referenced to the average of all electrodes, and separated back into individual time points. See table S2 for details on number of trials, channels and components removed.

***Table S1:*** *Total trials, number of channels removed, and number of components removed during cleaning of TEPs using pipeline one.*

| **Site, condition, time** | **Total trials** | **Channels removed** | **Components removed (ICA1)** | **Components removed (ICA2)** | **Components removed (SSP-SIR)** |
|---|---|---|---|---|---|
| PFC, DXM, Pre | 135.9 [124-145] | 0.8 [0-1] | 1.9 [1-5] | 10.4 [4-23] | 0.4 [0-2] |
| PFC, DXM, Post | 130.0 [95-144] | 0.7 [0-1] | 2.3 [1-5] | 10.6 [3-22] | 0.6 [0-4] |
| PFC, PBO, Pre | 137.8 [128-145] | 0.8 [0-1] | 2.0 [1-4] | 12.6 [3-27] | 0.4 [0-2] |
| PFC, PBO, Post | 136.6 [113-144] | 0.8 [0-1] | 1.9 [1-4] | 9.9 [2-19] | 0.4 [0-2] |
| PAR, DXM, Pre | 133.5 [115-142] | 0.9 [0-1] | 2.7 [1-4] | 11.3 [1-29] | 0.7 [0-4] |
| PAR, DXM, Post | 128.9 [92-145] | 0.8 [0-1] | 2.5 [0-4] | 14.9 [4-26] | 0.8 [0-3] |
| PAR, PBO, Pre | 132.9 [111-144] | 0.8 [0-1] | 2.6 [1-5] | 12.8 [3-28] | 0.6 [0-1] |
| PAR, PBO, Post | 139.6 [134-146] | 1.1 [0-2] | 2.7 [2-5] | 9.1 [3-21] | 1.5 [0-4] |

***NB:*** *Data are mean [range]. DXM, dextromethorphan; ICA, independent component analysis; PAR, parietal cortex; PBO, placebo; PFC, prefrontal cortex; SSP-SIR, signal-space-projection source-informed reconstruction; TEP, TMS-evoked potential.*

***Table S2:*** *Total trials, number of channels removed, and number of components removed during cleaning of TEPs using pipeline two.*

| Site, condition, time | Total trials | Channels removed | Components removed (ICA1) | Components removed (ICA2) |
|---|---|---|---|---|
| PFC, DXM, Pre | 134.6 [128-142] | 1.5 [0-5] | 0.9 [0-3] | 30.7 [24-38] |
| PFC, DXM, Post | 132.0 [92-141] | | | |
| PFC, PBO, Pre | 135.5 [129-140] | 1.8 [0-5] | 0.9 [0-3] | 29.7 [22-37] |
| PFC, PBO, Post | 133.2 [126-138] | | | |
| PAR, DXM, Pre | 134.1 [116-142] | 1.3 [0-3] | 2.1 [0-5] | 33.9 [24-44] |
| PAR, DXM, Post | 132.5 [92-140] | | | |
| PAR, PBO, Pre | 131.1 [97-140] | 1.6 [0-5] | 1.6 [0-4] | 32.1 [15-43] |
| PAR, PBO, Post | 135.2 [128-141] | | | |

***NB:*** *Data are mean [range]. Missing values indicate when data were concatenated and cleaning was applied equally between pre and post time points. DXM, dextromethorphan; ICA, independent component analysis; PAR, parietal cortex; PBO, placebo; PFC, prefrontal cortex; TEP, TMS-evoked potential.*

***Table S3:*** *Total segments, number of channels removed, and number of components removed during cleaning of resting EEG.*

| State, condition, time | Total segments | Channels removed | Components removed |
|---|---|---|---|
| Open, DXM, Pre | 124.4<br>[103-165] | 0<br>[0-0] | 21.8<br>[13-31] |
| Open, DXM, Post | 124.1<br>[114-139] | | |
| Open, PBO, Pre | 126.1<br>[85-162] | 0<br>[0-0] | 21.3<br>[11-29] |
| Open, PBO, Post | 122.5<br>[82-165] | | |
| Closed, DXM, Pre | 122.1<br>[110-147] | | |
| Closed, DXM, Post | 134.6<br>[103-197] | | |
| Closed, PBO, Pre | 121.8<br>[75-172] | | |
| Closed, PBO, Post | 125.8<br>[88-168] | | |

***NB:*** *Data are mean [range]. Missing values indicate when data were concatenated and cleaning was applied equally between pre and post time points and eyes open and eyes closed states. DXM, dextromethorphan; EEG, electroencephalography; PBO, placebo.*

### SUPPLEMENTARY RESULTS

*Effect of dextromethorphan on blood pressure*

There was a main effect of time on diastolic blood pressure ($F_{1,13}$=7.9, p=0.006), with pressure increasing across the experiment, however we could not detect any main effect of condition ($F_{1,13}$=0.2, p=0.71), or time × condition interaction ($F_{1,13}$=0.4, p=0.66). We could not detect any main effects of time ($F_{1,13}$=2.4, p=0.13) or condition ($F_{1,13}$=0.0, p=0.90), or a time × condition interaction ($F_{1,13}$=1.1, p=0.34) on systolic pressure, suggesting dextromethorphan and placebo did not have differential effects on blood pressure.

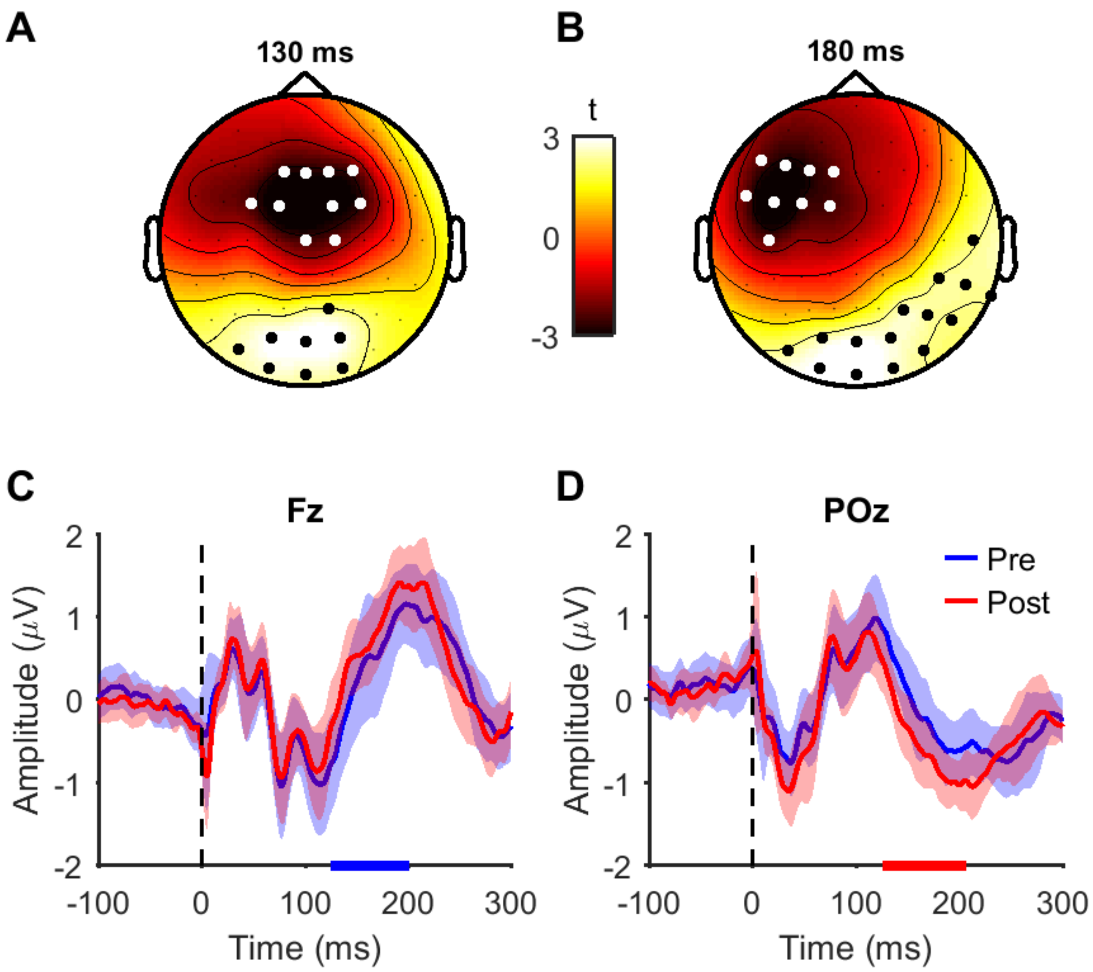


***Figure S1: Changes in TEP amplitudes following dextromethorphan after parietal cortex stimulation.*** *A-B) Topoplots displaying t-statistics at two time points during significant clusters indicating differences in TEP amplitudes pre and post dextromethorphan administration (positive cluster, p=0.006, 126-207 ms, black dots; negative cluster, p=0.0132, 125-201 ms, white dots). C-D) Plots from single electrodes in negative (C) and positive (D) clusters pre and post dextromethorphan administration. Shaded bars represent 95% confidence intervals. Solid bars on x axis indicate timing of significant negative (blue) and positive (red) clusters. Dashed black line indicates timing of the TMS pulse.*

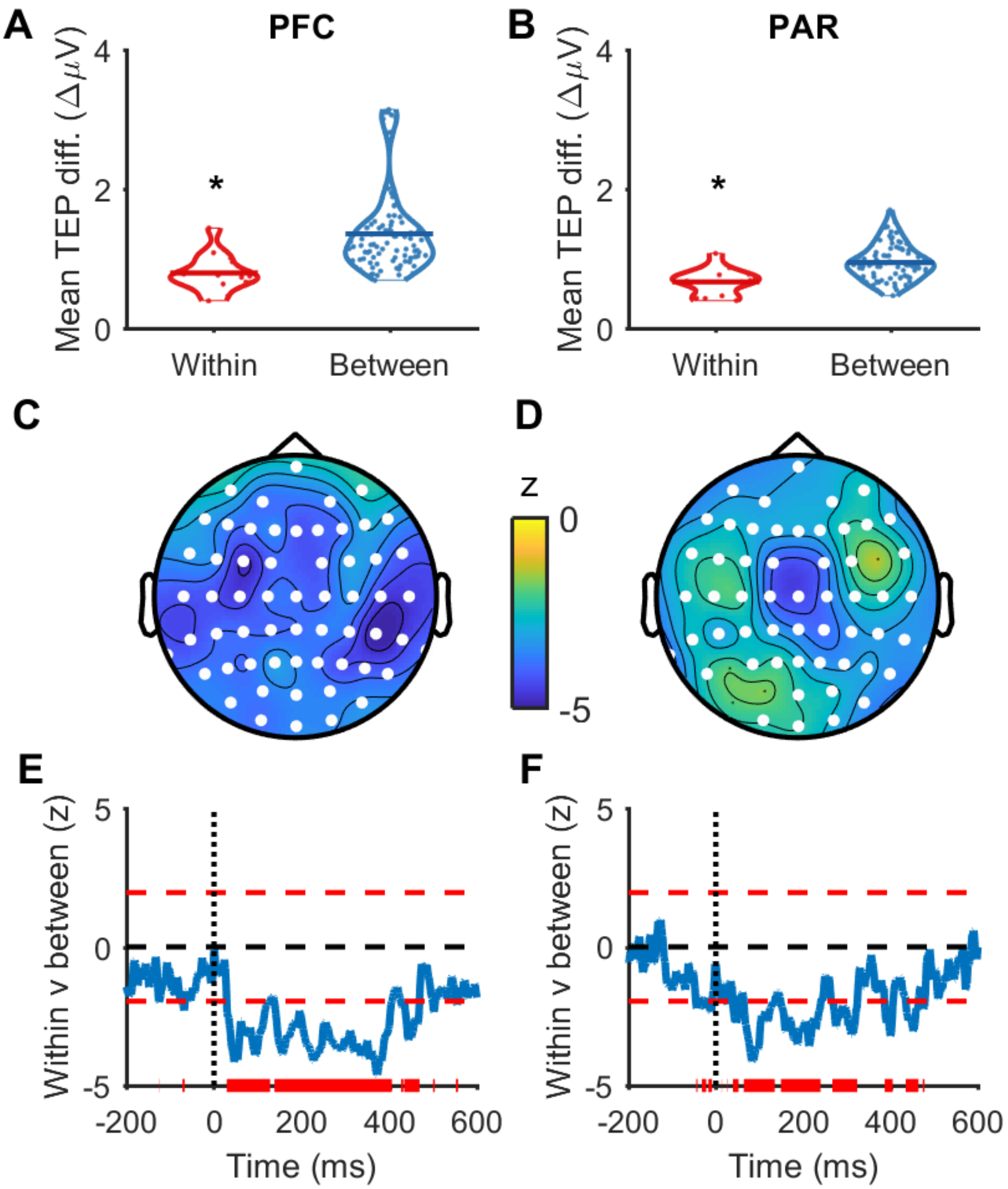


***Figure S2: Within- and between-subject variability in baseline TEPs across conditions using TMS-EEG cleaning pipeline two.*** *A-B) Mean absolute differences in baseline TEPs (15-500 ms, all electrodes) between the dextromethorphan and placebo condition within- and between-subjects following prefrontal cortex (PFC; A) and parietal cortex (PAR; B) stimulation. * indicates p<0.05 (Mann-Whitney U test). C-D) Topoplots displaying z-scores (Mann-Whitney U*

*tests) comparing within- and between-subject baseline TEP differences at individual electrodes (averaged across time between 15-500 ms) following PFC (C) and PAR (D) stimulation. Negative z-scores indicate within-subject TEP differences are less than between-subject TEP differences. White dots indicate $p<0.05$. E-F) Z-scores (Mann-Whitney U tests) comparing within- and between-subject TEP differences at individual time points (averaged across all electrodes) following PFC (E) and PAR (F) stimulation. Dotted black lines indicate the time of the TMS pulse. Dashed red lines indicate $z = \pm 1.96$. Solid red lines indicate $p<0.05$.*

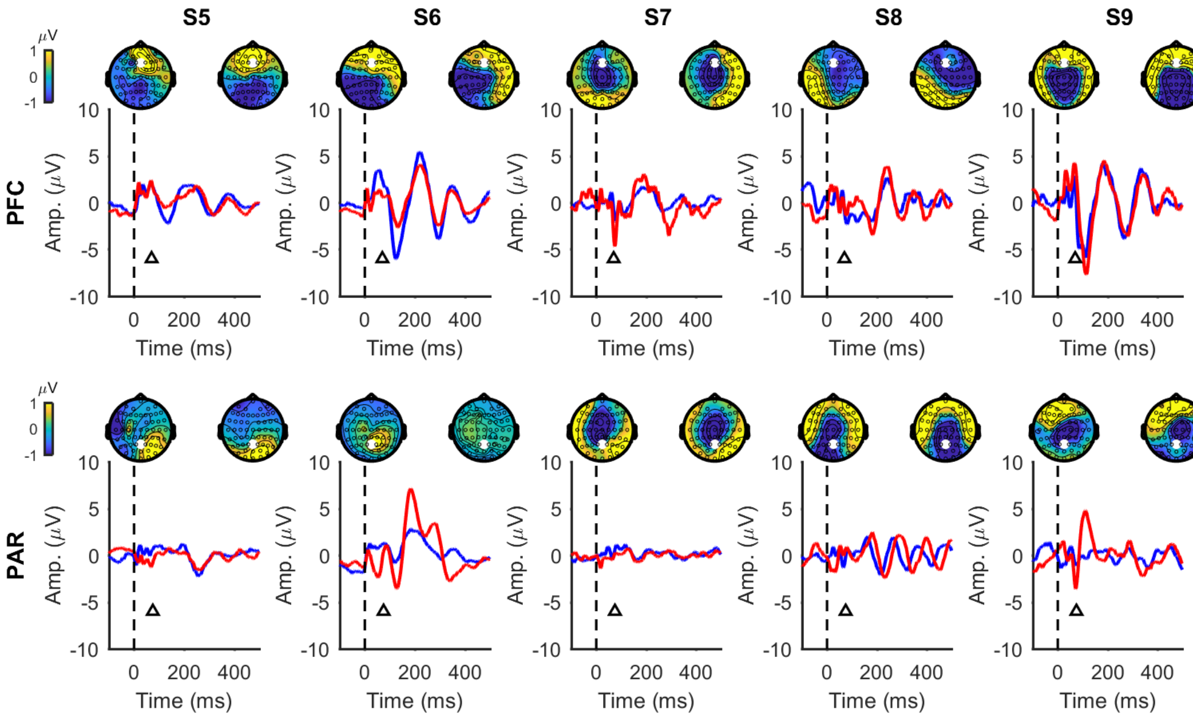


***Figure S3: Example of within- and between-subject variability in individual participants using TMS-EEG cleaning pipeline two.*** *The top row shows baseline TEPs from the dextromethorphan (blue lines, left topoplots) and placebo (red lines, right topoplots) sessions following PFC stimulation, and the bottom row following PAR stimulation, in five example participants (S5-S9). TEP line plots are taken from an electrode near the site of stimulation (indicated with white dot on topoplots; Fz for PFC stimulation; Pz for PAR stimulation), and TEP topoplots for a representative point in time (indicated with triangles; 70 ms for PFC; 75 ms for PAR). Both the shape and spatial distribution of the baseline TEPs are more similar within-subjects than they are between-subjects.*

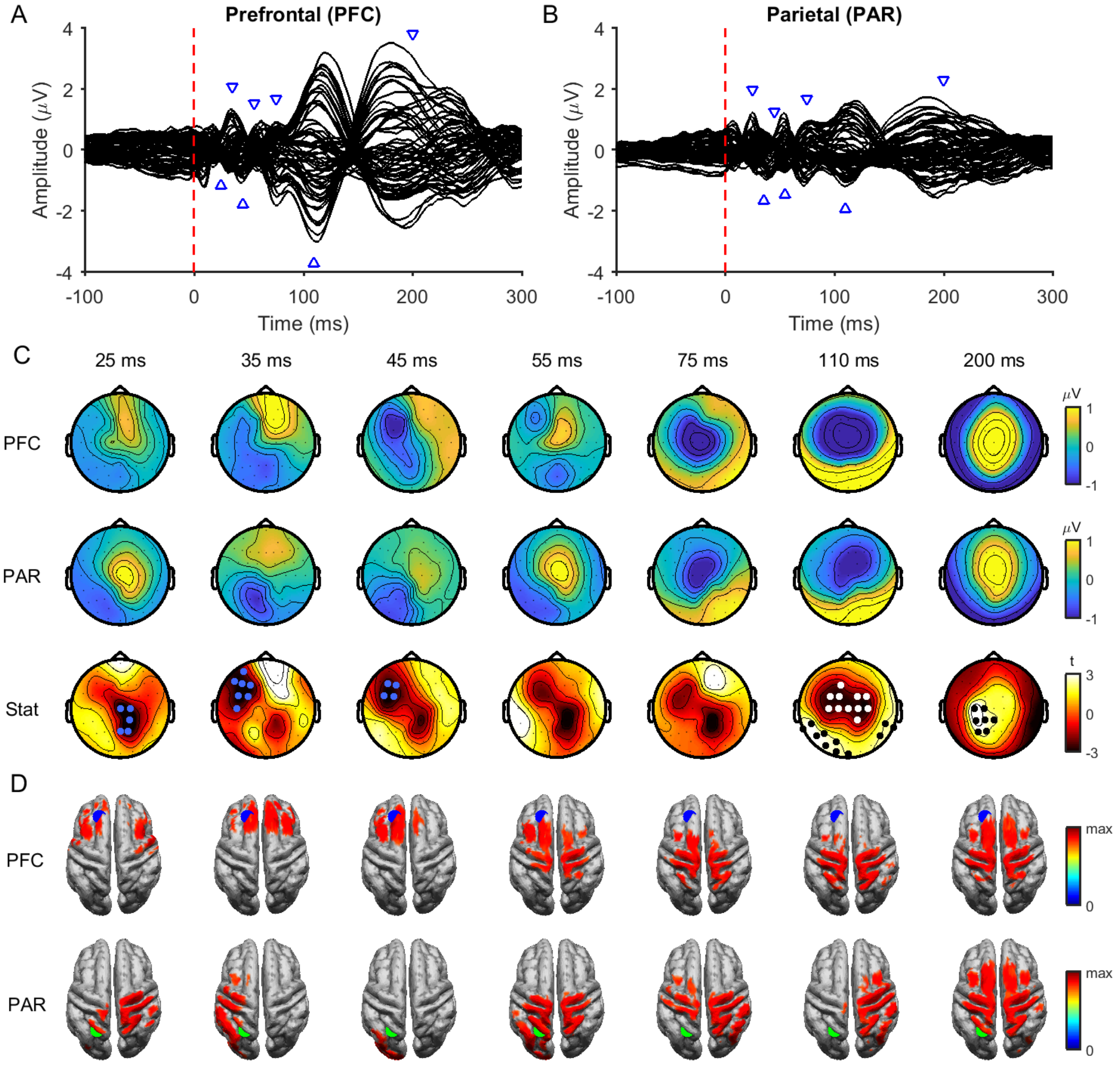


***Figure S4: Comparison of baseline TEPs between stimulation sites using TMS-EEG cleaning pipeline two.*** *Butterfly plots of grand average TEPs across all individuals following prefrontal (PFC; A) and parietal cortex (PAR; B) stimulation at baseline (averaged across conditions). The red dashed line represents the timing of the TMS pulse and the blue triangles the latencies plotted in C and D. C) Topoplots showing the grand average amplitude of TEPs at different time points following PFC (top row), and PAR stimulation (middle row). The bottom row shows t-statistics comparing the amplitude of PFC and PAR stimulation. White and black dots indicate significant negative and positive clusters (p<0.05; cluster-based permutation tests on*

*15-250 ms). Blue dots indicate significant clusters over shorter time windows (15-30 ms; 31-45 ms). D) Minimum-norm estimate source maps averaged across participants showing peak activity at each time point in C following PFC (top row) and PAR (bottom row) stimulation. Activity has been thresholded to 85% of maximum activity at each time point. The blue dot represents the target for PFC stimulation and the green dot the target for PAR stimulation.*

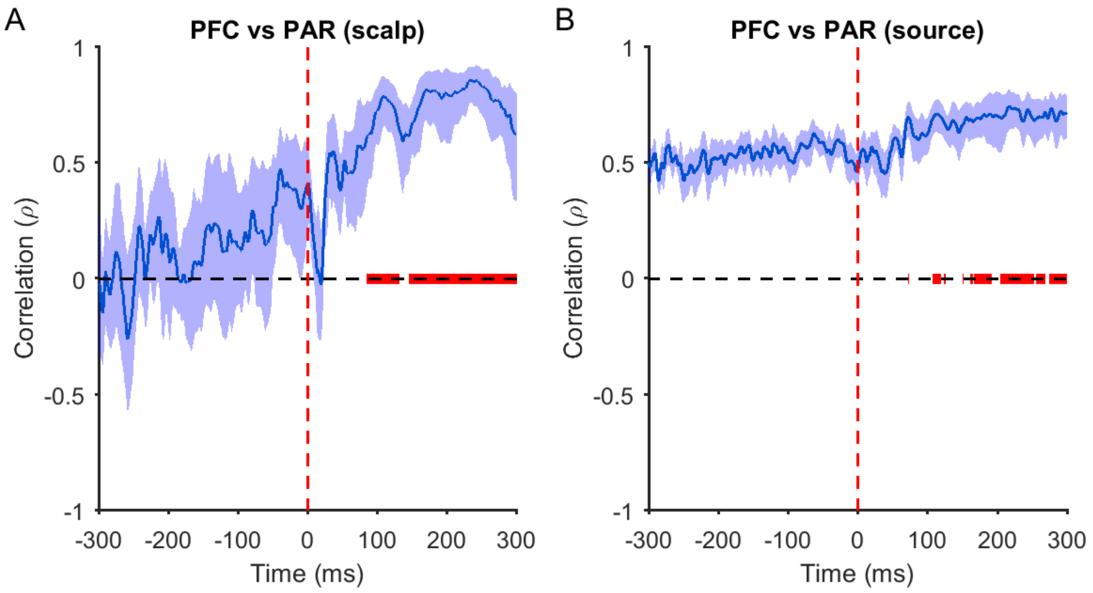


***Figure S5: Spatial correlations between prefrontal (PFC) and parietal (PAR) TEPs following TMS-EEG cleaning pipeline two.*** *Spearman correlations comparing the relationship between PFC and PAR TEPs at the scalp (A) and source (B) level for each time point. The thick blue line represents the mean rho values across individuals, and the shaded bars the 95% confidence intervals. The thick red line indicates post stimulation time points where correlations are greater than at equivalent pre stimulation time points (p<0.05; Mann-Whitney U test). Note that rho values were converted to z for statistics, then back to rho for plotting.*

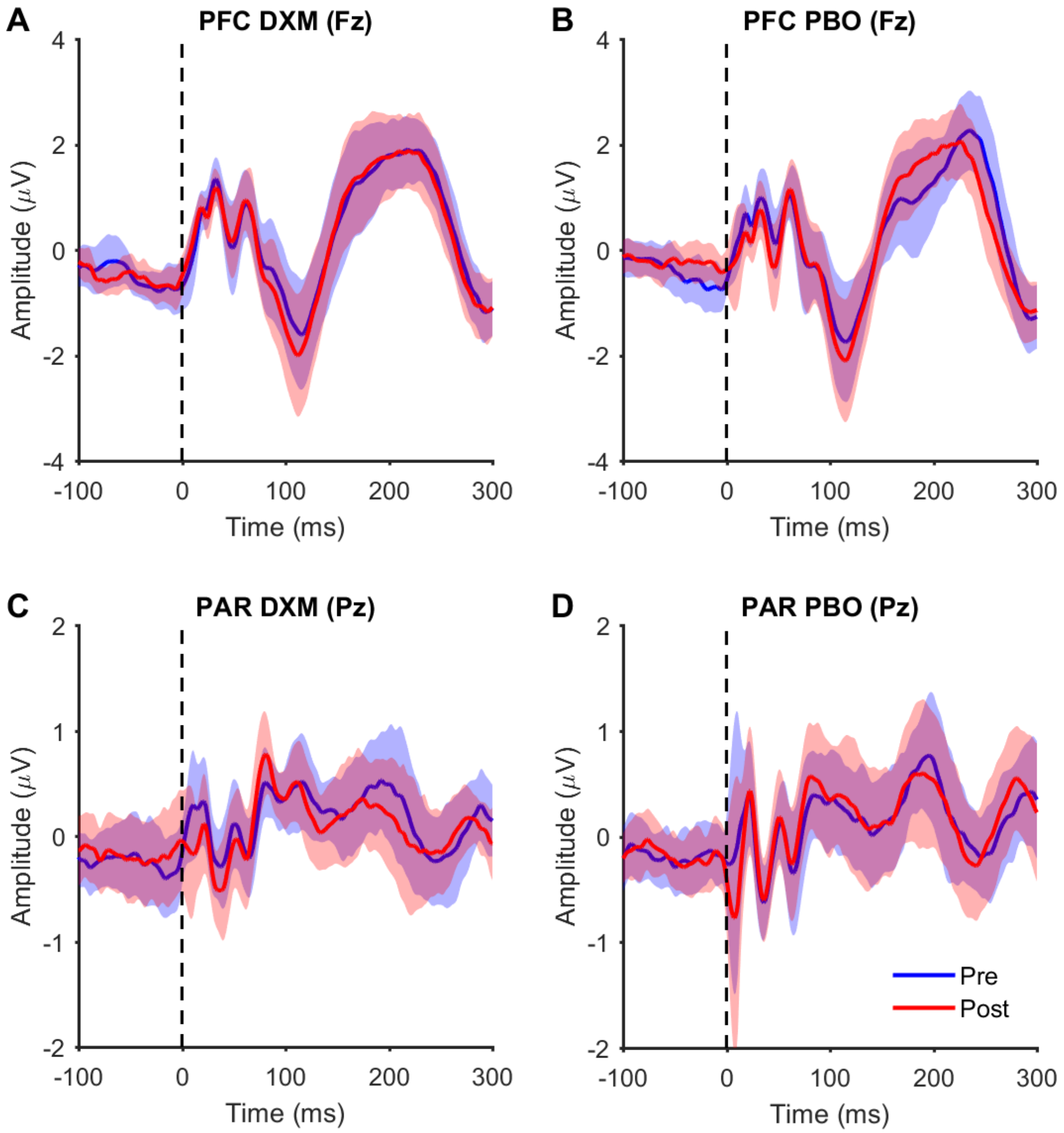


***Figure S6: TEPs from single electrodes following dextromethorphan (DXM) and placebo (PBO) using TMS-EEG cleaning pipeline two.*** *A-B) TEPs measured from the Fz electrode following prefrontal cortex (PFC) stimulation pre and post dextromethorphan and placebo administration. C-D) TEPs measured from the Pz electrode pre and post dextromethorphan and placebo administration. Thick coloured lines represent the group mean and shaded colour lines represent 95% confidence intervals.*

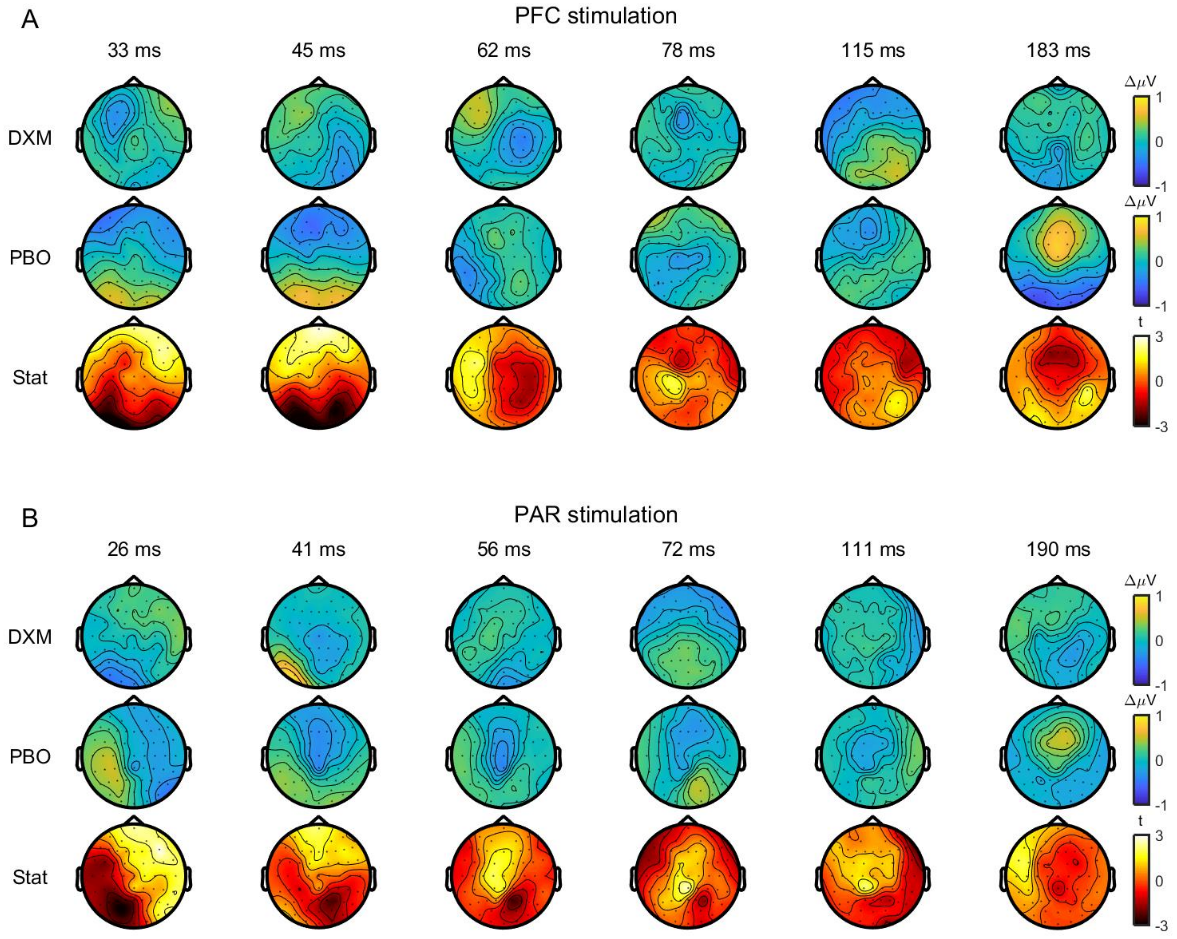


***Figure S7: Comparison of changes in TEPs following dextromethorphan (DXM) and placebo (PBO) using TMS-EEG cleaning pipeline two.*** *Topoplots showing changes in TEP amplitude at peak latencies following prefrontal (PFC; A) and parietal cortex (PAR; B) stimulation after dextromethorphan (top row) and placebo (middle row). Topoplots showing t-statistics (within-subject t-tests) comparing TEP changes between dextromethorphan and placebo are shown on the bottom row. No differences were observed between conditions (cluster-based permutation tests).*

***Table S4:*** *Distance from TMS target sites to best-fitting dipoles at baseline using pipeline two.*

| | **Distance from target (mm)** | **Distance from non-target (mm)** | **Goodness of fit (GoF)** | **p-value** |
|---|---|---|---|---|
| PFC (15-45 ms) | **54 [21-100]** | 70 [41-92] | 0.89 [0.80-0.96] | 0.046 |
| PFC (95-125 ms) | 94 [58-134] | **52 [15-89]** | 0.86 [0.65-0.96] | $2.16\times10^{-4}$ |
| PFC (175-205 ms) | 80 [41-128] | **52 [33-90]** | 0.85 [0.68-0.964] | $8.65\times10^{-4}$ |
| PAR (15-45 ms) | **48 [20-84]** | 87 [43-139] | 0.89 [0.65-0.95] | $2.59\times10^{-4}$ |
| PAR (95-125 ms) | **55 [24-88]** | 94 [78-114] | 0.89 [0.74-0.98] | $3.91\times10^{-5}$ |
| PAR (175-205 ms) | **46 [9-70]** | 80 [44-131] | 0.83 [0.64-0.98] | $8.55\times10^{-5}$ |

***NB:*** *Values in column 1-3 represent the mean [range]. Bold numbers indicate which site was closest to the best fitting dipole (target vs. non-target; p<0.05, Mann-Whitney U test). PFC, prefrontal cortex; PAR, parietal cortex.*

***Table S5:*** *Bayes factors comparing the change in TEP peak amplitude following dextromethorphan (DXM) vs. placebo (PBO) using pipeline two.*

| | **DXM vs PBO** | |
|---|---|---|
| **TEP peaks** | **PFC ($BF_{01}$)** | **PAR ($BF_{01}$)** |
| 33, 25 | 0.9 | 1.7 |
| 43, 41 | 0.9 | **3.5** |
| 60, 54 | **3.7** | 2.1 |
| 77, 73 | **3.4** | 1.2 |
| 115, 112 | **3.6** | 2.8 |
| 184, 194 | 2.8 | 2.8 |

***NB:*** *Values in column one represent the mean TEP peak latency for prefrontal cortex (PFC) and parietal cortex (PAR) stimulation respectively. Bold numbers indicate moderate evidence for no difference between conditions.*
